# Rapid resistance to pesticide control is predicted to evolve in an invasive fish

**DOI:** 10.1101/718593

**Authors:** Mark R. Christie, Maria S. Sepúlveda, Erin S. Dunlop

**Affiliations:** Department of Biological Sciences, Purdue University, 915 W. State St., West Lafayette, Indiana, 47907, USA; Department of Forestry and Natural Resources, Purdue University, 715 W. State St., West Lafayette, Indiana, 47907, USA; Aquatic Research and Monitoring Section, Ontario Ministry of Natural Resources and Forestry, Peterborough, Ontario, K9L 0G2, Canada; Environmental and Life Sciences Graduate Program, Trent University, Peterborough, Ontario, K9L 0G2, Canada

**Keywords:** eco-evolutionary dynamics, gene flow, invasive species, pesticide, rapid evolution, resistance, sea lamprey

## Abstract

Xenobiotic resistance is commonly found in species with short generation times such as bacteria, annual plants, and insects. Nevertheless, the fundamental evolutionary principles that govern the spread of resistance alleles hold true for species with longer generation times. One such example could occur with sea lamprey (*Petromyzon marinus*), a parasitic invasive species in the Laurentian Great Lakes that decimated native fish populations prior to its control with the pesticide 3-trifluoromethyl-4-nitrophenol (TFM). Since the 1950s, tributaries have been treated annually with TFM, where treatments effectively remove most, but not all, larval sea lamprey. We developed an eco-genetic model of sea lamprey to examine factors affecting the evolution of resistance and found that resistance alleles rapidly rise to fixation after 40-80 years of treatment, despite the species’ relatively long generation time (4-7 years). The absence of natal homing allows resistant individuals to spread quickly throughout the entire system, but also makes the early detection of resistance challenging. High costs of resistance and density independent reproduction can delay, but not prevent, the onset of resistance. These results illustrate that sea lamprey have the potential to evolve resistance to their primary control agent in the near future, highlighting the urgent need for alternative controls.

## Introduction

The rapid evolution of resistance to xenobiotic compounds has been widely documented in microbes^1,2^, fungi^3^, invertebrates^4,5^, and plants^6,7^. Resistance has also been documented in vertebrates^8–10^, but has largely been constrained to taxa with high fecundity and short generation times. Nevertheless, the fundamental principles governing the evolution of resistance still apply to species with longer generation times. If the selection pressure imposed by a xenobiotic is strong, wide-spread, and consistently applied year after year, then resistance may still evolve. The need to control nuisance pest species through various means including xenobiotics continues to rise with globalized trade and climate change introducing species into new habitats. Aquatic species are no exception, with some of the more recent threats including invasive fishes^11,12^. Invasive species cost billions of dollars to control and they can impact ecosystems in drastic ways^13^. Thus, there is a need to consider the pace at which a broader range of taxa, including those with longer generation times such as fishes, could evolve resistance to control measures.

In this study, we examine the timelines of pesticide resistance evolution in an aquatic invasive fish species, the sea lamprey (*Petromyzon marinus*). In the Laurentian Great Lakes, a powerful chemical pesticide has been applied to control invasive sea lamprey for over 60 years^14^. The pesticide, 3-trifluoromethyl-4-nitrophenol (hereafter TFM), has been applied on an annual basis since the 1950s to Great Lakes tributaries, targeting sea lamprey during their larval stage. Because the pesticide kills most, but not all, larval sea lamprey^15,16^, some individuals are likely to survive exposure. If this process is repeated over a long enough period, then resistant individuals could increase in frequency – a scenario documented in many systems where pests have been controlled by chemical means^17^. The evolution of resistance would greatly reduce the effectiveness of TFM, which remains the primary control agent, and could result in large declines to native fish populations due to increases in sea lamprey abundance. These declines in fish abundance would not only have large ecological repercussions, but could also result in billions of dollars in economic damage^18,19^.

As adults, sea lamprey feed on the blood and tissues of host species, including species of commercial importance such as salmon and lake trout. Invasive sea lamprey impact native fish populations by wounding and often killing the host fishes that they parasitize^18,20^. Following their invasion in the late 1930s, sea lamprey contributed to the catastrophic loss of economically valuable commercial and recreational fisheries in Canada and the United States^21^. In response to the proliferation of sea lamprey throughout the Great Lakes, there was an immediate and concerted effort to develop efficient means of control. One effort, initiated in the 1950s, involved testing over 6,600 chemical compounds on sea lamprey and other fish species^22^. The organic compound TFM was found to effectively kill larval sea lamprey and had few detectable effects on other fish species at low concentrations. TFM control was deemed highly successful and sea lamprey abundance was reported to have been reduced by approximately 90%, resulting in recovery of economically important fisheries^23^. This outcome is one of the few documented cases of an invasive vertebrate species being successfully controlled at an ecosystem scale by a pesticide. Alternative control measures have been implemented on smaller scales, but none have proven as effective as TFM, with the possible exception of existing barriers that prevent upstream migration to spawning habitat^24–26^. However, barriers are not present on most tributaries and there is increased pressure from stakeholders to remove existing barriers in order to restore natural connectivity in waterways^27^. Thus, there is a continued reliance on TFM to manage invasive sea lamprey populations in the Great Lakes. This reliance on a single control measure can be risky; only a handful of manufacturers produce TFM and thus both prices and supplies of TFM could change. More importantly, reliance on a single chemical control measure has been shown to increase the chances of resistance evolution^28,29^.

There are several reasons why invasive sea lamprey could be evolving resistance to TFM. First, sea lamprey, like many pest species, have high fecundity. A single female lamprey can produce over 100,000 eggs in the Great Lakes^30^ and substantially more in its native range^31^. With such high fecundity, it may not take long for novel mutations that confer resistance to appear in sea lamprey offspring. Alternatively, there may be resistant individuals already present within the population (i.e., selection may act on standing genetic variation). Second, the selection pressure imposed by TFM may be substantial. Similar to many plant and insect pests, sea lamprey larvae can be found at high density; sea lamprey larvae congregate in lentic, sandy tributaries and this life-history strategy means that large numbers of individuals can be targeted with a single application of TFM. Estimates of larval mortality per stream treated with TFM range from 80-99% ^32^, representing a substantial selective event. Third, the selection pressure from TFM has been present for nearly 60 years. TFM has been applied to Great Lakes tributaries almost every year since 1958^14^. Thus, the combination of high fecundity, strong selection, and long-term treatment make the evolution of resistance a distinct possibility.

Despite the possibility for the evolution of resistance, there are several reasons why resistance evolution may be prevented or delayed. One reason is that sea lamprey have overlapping generations and they can live for a relatively long time (e.g., up to 19 years in its native range^33^). This longevity means that sea lamprey generation times are longer and their response to selection slower in real time in comparison with many other pest species. Assuming an average generation time of between 3 and 5 years, invasive sea lamprey may have been exposed to TFM for 20-30 generations, which may not be sufficient time for resistance to evolve (but see ^34–36^). Another reason that resistance may not evolve is the high rate of dispersal and subsequent gene flow found in this species. Adult sea lamprey do not return to their natal streams to spawn, instead choosing spawning sites via the detection of pheromones released by larvae^37^. Streams with more larvae release greater pheromone concentrations and attract more adults to spawn. This mixing of adults during reproduction results in high gene flow and near-panmixia throughout the Great Lakes^38^. The effects of this lack of natal philopatry on the evolution of resistance remain largely unknown, but the high gene flow between treated and non-treated streams may allow for reservoir populations to limit the spread of resistant genotypes^39^. Previous empirical work has not found any evidence of TFM resistance in invasive sea lamprey^34^. However, there were limitations to the empirical analysis and, as we show in this manuscript, detection of resistance in anything but its latest stages requires vast sample sizes.

In this study, we used the sea lamprey as a model system to examine the likelihood and timelines of resistance evolution in a fish species with a relatively long generation time and that has experienced a potentially potent selective pressure from the persistent application of a chemical pesticide. To predict the evolution of resistance in this system, we built an eco-genetic model^40^ that includes important aspects of sea lamprey life history, spatial structure, and TFM control. Specifically, we answer five questions: 1. How quickly can resistance evolve under varied strengths of selection? 2. What level of treatment intensity results in the fewest lamprey over the long term? 3. How does the lack of natal philopatry affect the development of resistance? 4. How do costs of resistance affect the persistence and timing of resistance? and 5. How does the relative magnitude and timing of density dependent and density independent process affect resistance? We conclude with a discussion of caveats and model assumptions, but nevertheless suggest that if agencies are as effective at controlling invasive lamprey as they believe, then the evolution of resistance is imminent.

## Materials and Methods

To examine the evolution of TFM resistance in sea lamprey, we constructed a forward-time, individual-based model (IBM). The model is a type of eco-genetic model^40^, which follows the evolution of key traits while also including important ecological feedbacks such as density-dependent reproduction and survival. The model is fully age and stage structured (including larvae, parasitic juveniles, and reproductively mature adults) and is based on life history characteristics of sea lamprey from the Great Lakes (see Supporting Information for detailed life history information; see Table 1 for default parameters). Each simulation was initiated by creating larval and juvenile individuals of varying ages and sexes (50:50 sex ratio). Larvae were split among tributaries and juveniles were assigned to a single parasitic population^20,38^. All individuals within the model were assigned a growth parameter, *k*, as a random deviate from a normal distribution with a mean of 0.001 and a variance of 0.0002. This growth parameter remained with the individual for their lifetime and was used in the Von Bertalanffy growth equation to determine size at age^41^:

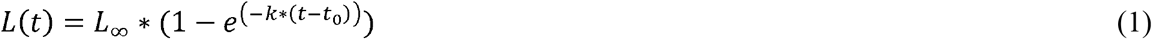

where *L*(*t*) equals the body length of an individual at time *t*, *L*_∞_ equals the maximum length, here set to 150 mm for larvae and 800 mm for juveniles^42^, *k* equals the individual growth parameter described above, *t* equals the individual’s age in days, and where to was set *t*_0_ −15. Examples of individual growth curves for larvae and juveniles using these parameters are illustrated in Figure 1. Size-at-age was used to determine when individuals would transition between stages. When a larva was greater than or equal to 120 mm in length, the individual transformed into a parasitic juvenile^15,32^. Likewise, when a parasitic juvenile was greater than or equal to 450 mm in length, the individual transitioned to reproductive maturity^42,43^.

**Table 1:** Model parameters and the default and range of values reported.

| Parameter description | Default value | Tested values |
| --- | --- | --- |
| Number of years | 200 | -- |
| First year of TFM treatment | 50 | -- |
| Year resistant adult added | 70 | 50, 70, 90 |
| TFM mortality rate | 0.9 | -- |
| Number of tributaries treated each year | 0.2 | 0.05, 0.1, 0.15, 0.2, 0.25, 0.3, 0.35, 0.4 |
| Gene flow | No homing | IBD: 0.01, 0.05, 0.10, 0.20 |
| Resistance cost | 0 | 0, 0.1, 0.2, 0.3, 0.4, 0.5 |
| Number of larval populations/tributaries | 20 | 20, 40, 60 |
| Larval carrying capacity per tributary | 1200 | 1200, 12000 |
| Juvenile carrying capacity | 8000 | 8000, 80000 |
| Density dependent/independent reproduction | 50 | 50, 25, 10, 9, 8, 7, 6, 5, 4, 3, 2 |

**Figure 1.**
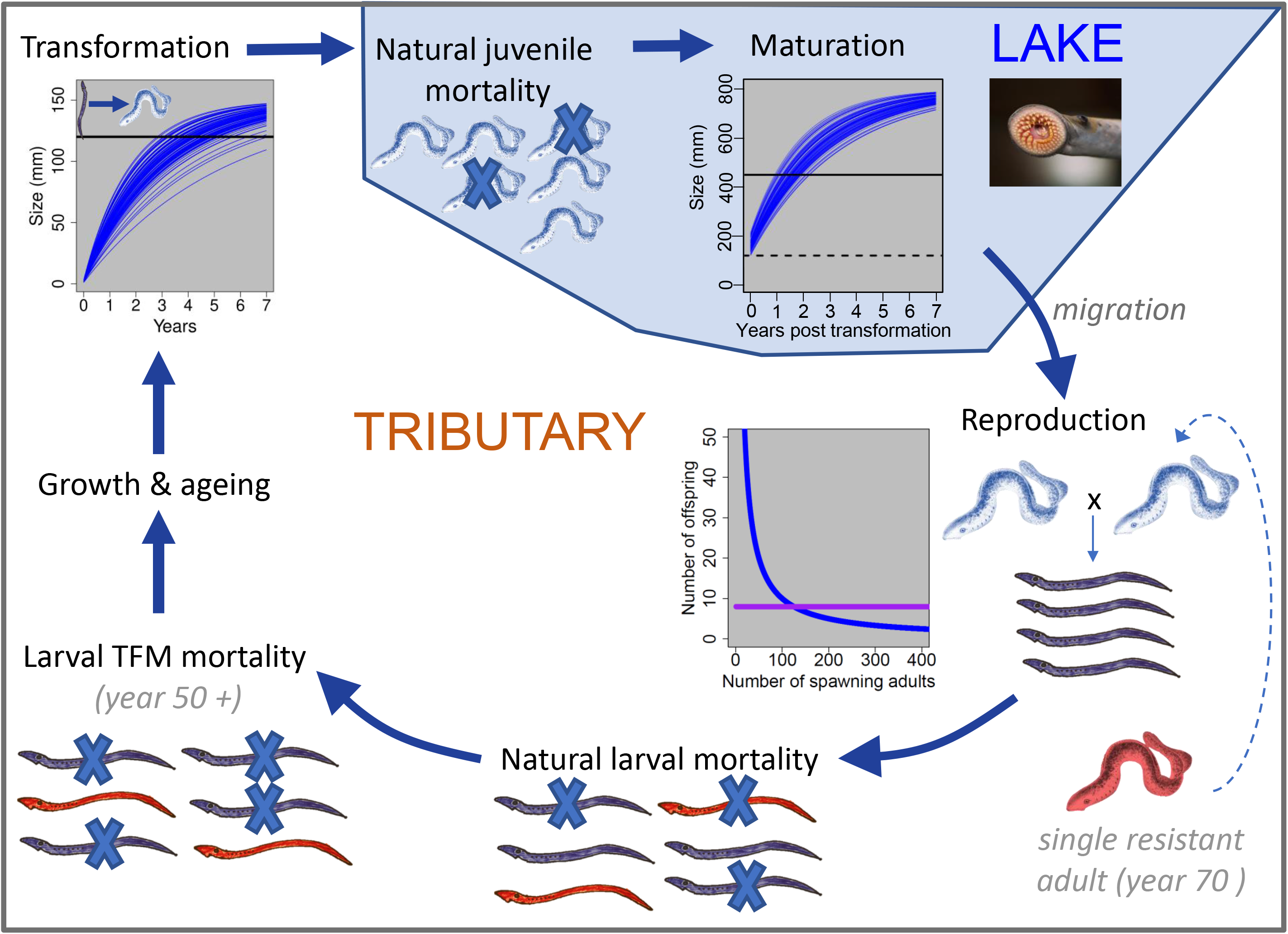
Illustration of the eco-genetic model of sea lamprey. Starting with growth and ageing, larval sea lamprey undergo metamorphosis between the larval ammocoete stage and the parasitic juvenile stage based on size-at-age. Each individual has a unique growth curve, with transformation occurring between ages 2-7 years after a size threshold is met (see inset where each blue line represents a different individual through time). In the lake, natural juvenile mortality is followed by maturation, which occurs after a second size threshold is surpassed. After migrating to tributaries, adults form pairs and reproduce. The number of offspring produced per spawning pair varies from density dependent (blue line, default) to density independent (purple line). After reproduction, adults die, and natural larval mortality occurs to reduce each larval tributary back to its carrying capacity. Starting in year 50, additional larval mortality occurs via treatment with the pesticide TFM. A single resistant adult is introduced into a randomly selected tributary at year 70. Resistant individuals are assumed to survive treatment with TFM. Model steps are repeated for a total of 200 years.

The first step in the model was ageing, where every individual was aged by one year (Figure 1). This step was followed by transformation (*i.e.*, larvae to juveniles) or maturation (*i.e.*, juveniles to adults) as dictated by size-at-age. After transformation, new juveniles were added to the single, panmictic lake-inhabiting population. After maturation, adults were assigned back to larval populations (*i.e*., tributaries) based on the relative abundances of larvae currently inhabiting each tributary. That is, in accordance with their life history, more adults were assigned back to large larval populations than to small larval populations with the following equation:

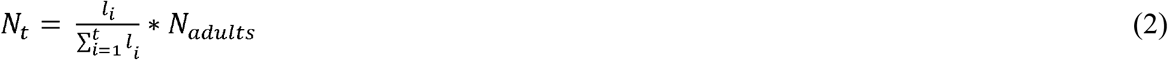

where N_t_ equals the total number of adults assigned back to tributary *t*, *l*_i_ equals the total number of larvae in tributary *t* and *N*_adults_ equals the total number of adults ready to spawn. This step was implemented to mimic the observation that adult sea lamprey, unlike salmon and many other anadromous fishes, do not return to their natal streams (see Supporting Information for detailed life history information). After randomly allocating adults to tributaries based on larval population sizes, males and females were paired and allowed to reproduce. Reproduction was modeled as both density dependent and density independent (though the default was density dependent; Table 1, Figure 1) and the number of offspring per pair was set as:

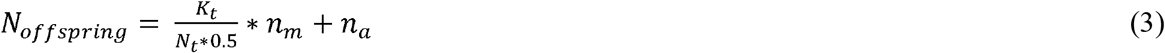

where *K_t_* equals the tributary specific carrying capacity, *N_t_* was calculated with equation (2) and *n_m_* and *n_a_* are constants that control the strength of density dependence. Previous empirical work in this system has shown that Great Lakes sea lamprey have highly compensatory population dynamics^44^, and so our default parameters assumed that egg and larval survival through the first year was also density dependent (Table 1). During reproduction each pair created *N*_*offspring*_ with each offspring inheriting, in Mendelian fashion, one randomly selected allele from each of their parents to create genotypes that determine resistance status (see details below). Variance in the number of offspring per pair was introduced with subsequent larval mortality. After reproduction, all adults were removed from the model simulating the strict semelparity found in adult sea lamprey.

Modeling larval mortality was the next step. This natural mortality occurred regardless of resistance status. For each tributary, a random deviate from a normal distribution was used to determine the number of surviving larvae. The mean of the normal distribution was equal to the carrying capacity for that particular tributary and the default standard deviation was set to 70 allowing for both density dependent and density independent mortality. Juvenile mortality was procedurally identical to larval mortality except that mortality was applied to the juvenile (parasitic, lake-inhabiting) population (substituting a lake-wide carrying capacity, Table 1). The steps (growth, transformation, maturation, reproduction, and mortality) were repeated for the first 50 years of each simulation to allow the age and stage structure to fully develop. At year 50, TFM was applied after the natural larval mortality step (Figure 1). A parameter within the model allowed us to set the number of tributaries to treat with TFM each year. Throughout the Great Lakes, larval abundance surveys are conducted annually, and the tributaries are rank ordered with the tributaries found with highest lamprey abundance given priority for treatment^14^. Similarly, in our model we rank ordered each population by the number of larvae and treated the top ranked tributaries. For example, if we set the number of treated tributaries per year to 5, we would treat the 5 largest larval populations. We assumed that TFM mortality was 0.9^34^, meaning that 90% of susceptible larvae within a tributary would die after exposure to TFM. The combination of both the number of treated tributaries and the TFM mortality rate determined the total number of larval lamprey killed by TFM each year.

Resistance was modeled as a single locus trait as small numbers of loci of large effect have dominated resistance studies across diverse taxa^45^. Here, resistance was modeled as a dominant trait; individuals heterozygous for the resistance allele were also conferred full resistance though they only passed on the resistance allele to half of their offspring. There were no resistant individuals at the beginning of each simulation. At year 70, a single resistant adult was added to the population of juveniles that had just transitioned to the adult stage. The resistant individual was added 20 years after the start of TFM to allow us to disentangle the effects of treatment and resistance, but the actual year a resistant adult was added had no effect on the results (e.g., resistance developed *x* years after the introduction of a resistant adult regardless of whether it was added at year 50, 70, or 90). We also added a parameter to examine costs associated with resistance, which is often found in empirical studies of resistance^19^. Cost of resistance was modeled as a decrease in reproductive success. For example, if the parameter was set to 0.1, then a resistant individual would have a 10% reduction in reproductive success (here, set as the number of offspring) relative to a non-resistant individual.

### Parameter exploration

To test the effect of TFM treatment on the development of resistance we first varied the number of treated tributaries holding all other parameters constant (default values; Table 1). We examined the effects of applying TFM each year to 5, 10, 20, and 30% of all tributaries. For each year, we measured the number of parasitic juveniles and the number of resistant larvae. To ensure that the results scaled with the larger absolute population sizes observed in the field, we re-ran these analyses setting larval and juvenile carrying capacities to be an order of magnitude larger than the default vales (Table 1). We next examined the effects of natal homing on the spread of resistance using the default parameters (Table 1) except for adult migration. We first examined a linear stepping stone model of migration^46^ and varied gene flow of adults among natal tributaries from 1 to 20% per year. This alternative stepping stone scenario was run in addition to the default no-homing migration pattern unique to sea lamprey. We kept track of the number and geographic spread of resistant larvae and, for each level of gene flow, measured the proportion of 100 replicates that had at least one resistant larva at model year 200.

We next examined the costs of resistance by modeling three costs of resistance (0, 0.1, and 0.4 reductions in fitness) and stopping treatment at year 100 (30 years after the introduction of a resistant adult). By stopping treatment, we could directly examine how quickly resistant individuals were removed from the population after the survival advantage associated with resistance was removed. To explore costs of resistance in more detail, we varied the cost of resistance from 0 to 0.5 in intervals of 0.1 and measured the proportion of resistant larvae as a function of sample year. We first assumed a perfect detection probability (i.e., all resistant lamprey were successfully identified), but later relaxed this assumption to examine the relationship between sampling effort (% of total larvae sampled), cost of resistance, and probability of detection. Lastly, we varied the strength of density-dependent reproduction by varying the parameter *n_m_* in equation 3 (see Supporting Information for details). The model was developed in R 3.5.0^47^ and run on a high-performance computing cluster. For every unique combination of parameters, 100 independent simulations were performed.

## Results

Treating tributaries with TFM effectively reduces the number of parasitic juveniles found in the system, however, this effectiveness is directly tied to the number of larval populations treated each year. When only 5% of tributaries are treated each year, there is only a small reduction in the number of parasitic juveniles (Figure 2*a*). Conversely, when 30% of the tributaries are treated with TFM each year there was an 81% reduction in the total number of parasitic juveniles (Figure 2*g*). This result was consistent regardless of the total number of individuals or tributaries included in the model (Figure S1; Figure S2). We also found that the total number of resistant larvae at year 200 (the last year in our model simulations) varied with the total number of treated tributaries (Figure 2*b,d,f,h*). Under the high selection intensity of 30% of tributaries treated, resistance took an average of 30 years to develop. Under more moderate selection intensities, resistance took between 51 and 109 years to develop (Figure 2*d, f*).

**Figure 2.**
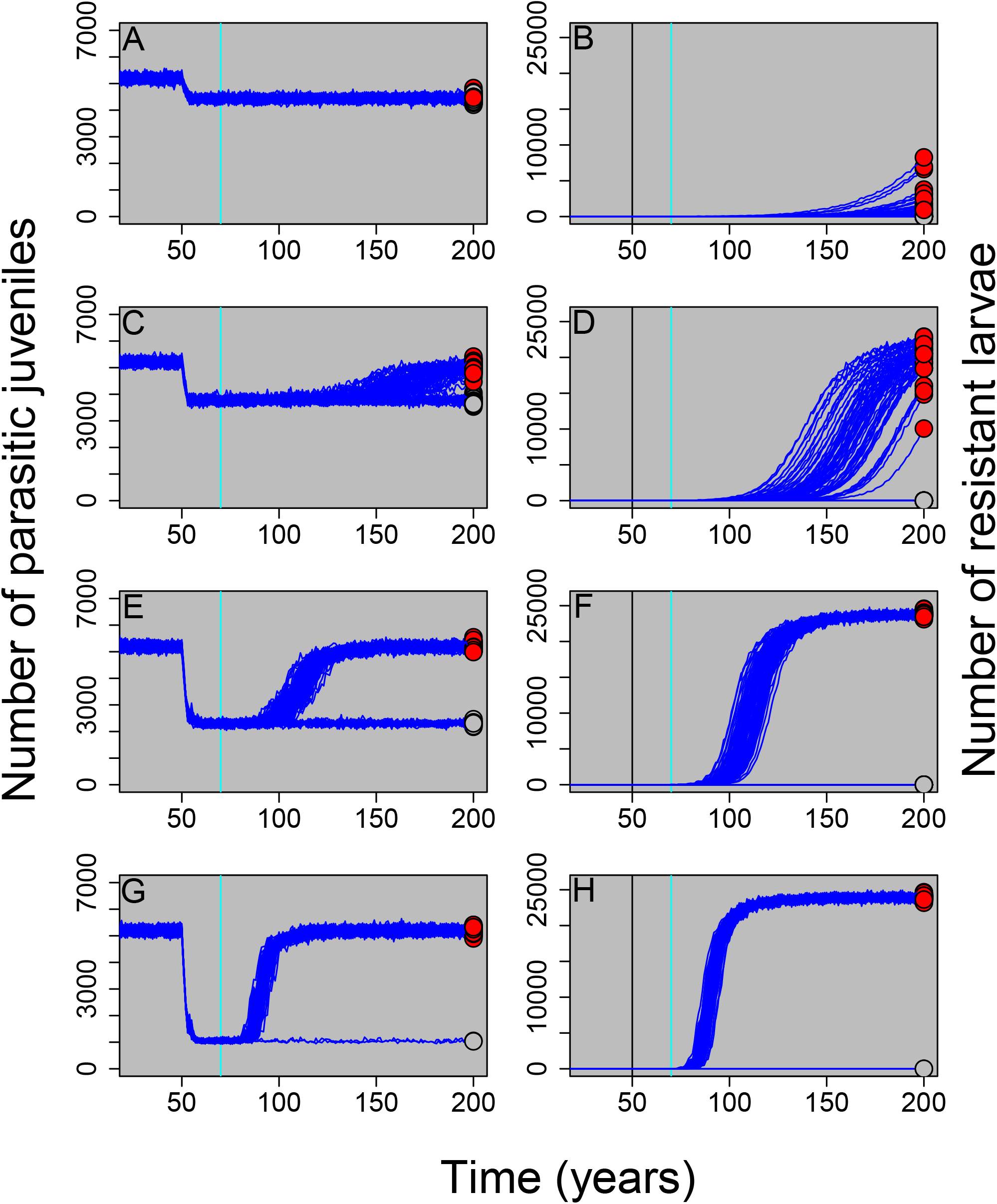
Parasitic juvenile and resistant larval abundances through time where the proportion of tributaries treated each year equaled 0.05 (panels A, B), 0.1 (panels C, D), 0.2 (panels E, F), and 0.3 (panels G, H), respectively. The number of parasitic juveniles includes both resistant and non-resistant juveniles (panels A,C,E,G) whereas the number of resistant larvae does not include non-resistant larvae (panels B,D,F,H). TFM treatment was started at year 50 (vertical black line) and a single resistant adult was introduced in year 70 (vertical blue line). Individual simulation results for one hundred replicates are shown for each scenario. At year 200, red circles represent populations with at least one resistant individual and grey circles represent populations with no resistant individuals.

The proportion of simulations that did not develop resistance was higher when the number of treated tributaries was lower (Figure 3). In other words, when selection intensity was low, the likelihood of successful reproduction by the single resistant adult introduced at year 70 was reduced (Figure 3). When resistance does evolve, the total (i.e., cumulative) number of parasites in the system over time is equivalent between heavily-treated systems where resistance often evolves and lightly-treated systems where resistance does not evolve as often (Figure 3). Combined, these results illustrate that a low selection intensity regime may result in the fewest parasitic lamprey over the long term because: 1. resistance is less likely to evolve (Figure 3*a*), and 2. when resistance does evolve it takes a much longer time for the number of resistant larvae to increase in frequency (Figure 2*b*). Thus, while a high selection intensity treatment regime will result in fewer parasitic lamprey over the short term, a low selection regime can ultimately result in fewer total lamprey in the system over the long term.

**Figure 3.**
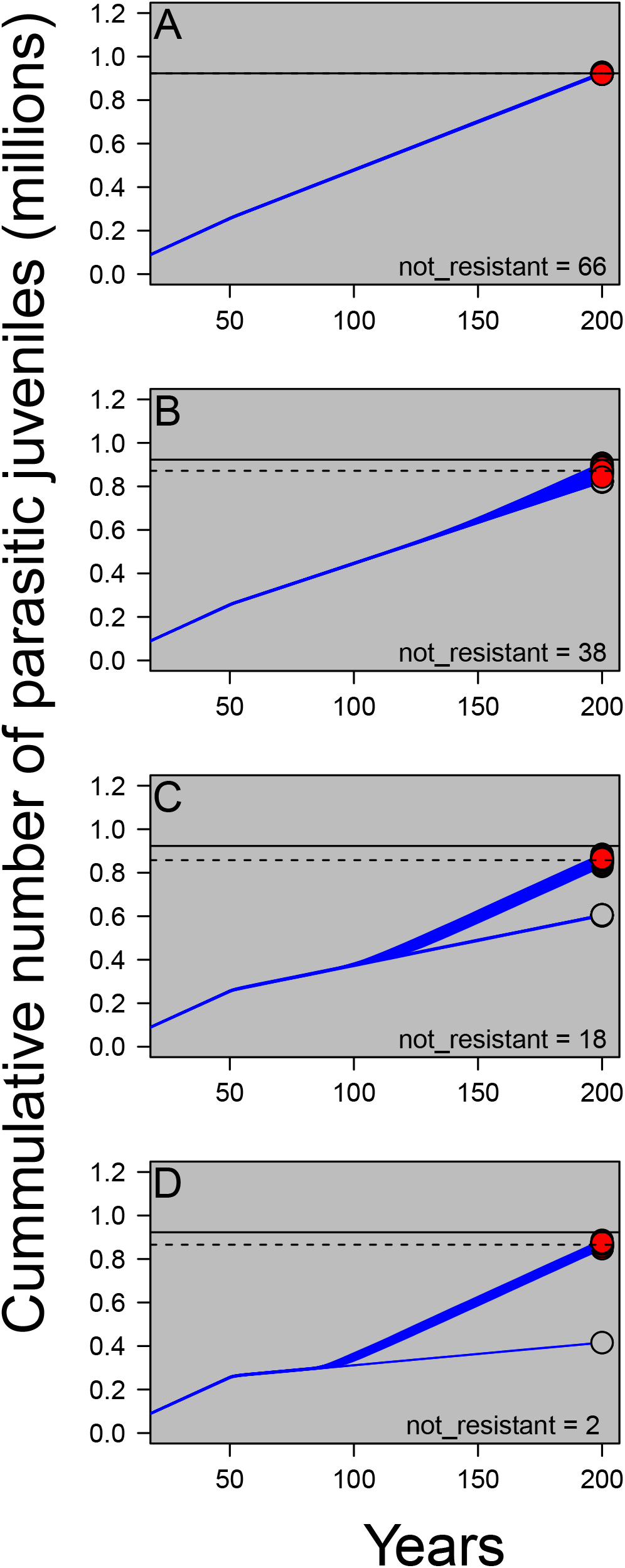
Cumulative parasitic juvenile abundances through time where the proportion of tributaries treated each year equaled 0.05 (panel A), 0.1 (panel B), 0.2 (panel C), and 0.3 (panel D), respectively. TFM treatment was started at year 50 and a single resistant adult was introduced in year 70. One hundred replicates (dark blue lines) were run for each scenario. Red circles represent populations with at least one resistant individual at year 200 and grey circles represent populations with no resistant individuals. The total number of simulations (out of 100) where no resistant individuals survived to year 200 are reported in the bottom right of each panel (“not resistant”). The solid horizontal line equals the mean number of parasites at year 200 where 5% of tributaries were treated each year (plotted on all panels, but measured from panel A). The dashed horizontal line represents the mean number of parasites at year 200 for each additional treatment intensity. Notice that when resistance evolves, higher treatment levels of TFM treatment do not substantially decrease the total number of parasites in the system over the long run. If resistance never evolves (grey circles), TFM will continue to be an effective management tool.

We also found that the sea lamprey life history strategy for identifying spawning habitat by cuing in on larval abundance greatly increases both the speed at which resistant individuals spread throughout the system and the proportion of simulations where resistance developed. For example, 15 years after the introduction of a resistant adult, resistant larvae had spread to 70% of larval tributaries when sea lamprey adult abundance was dictated by larval abundance (Figure 4*c*). By contrast, if larvae followed an isolation-by-distance pattern of dispersal, then only 5% of larval populations would be colonized 15 years after the introduction of a resistant adult (Figure 4*b*). Furthermore, even if gene flow among neighboring populations was as high as 20% per year (which would be high for most vertebrates), the proportion of simulations where resistance developed remained considerably lower than the actual migration strategy employed by lamprey (48% vs. 82%; Figure 4*d*). These results illustrate that the lack of natal philopatry in invasive sea lamprey greatly increase the probability of resistance spreading throughout the entire system.

**Figure 4.**
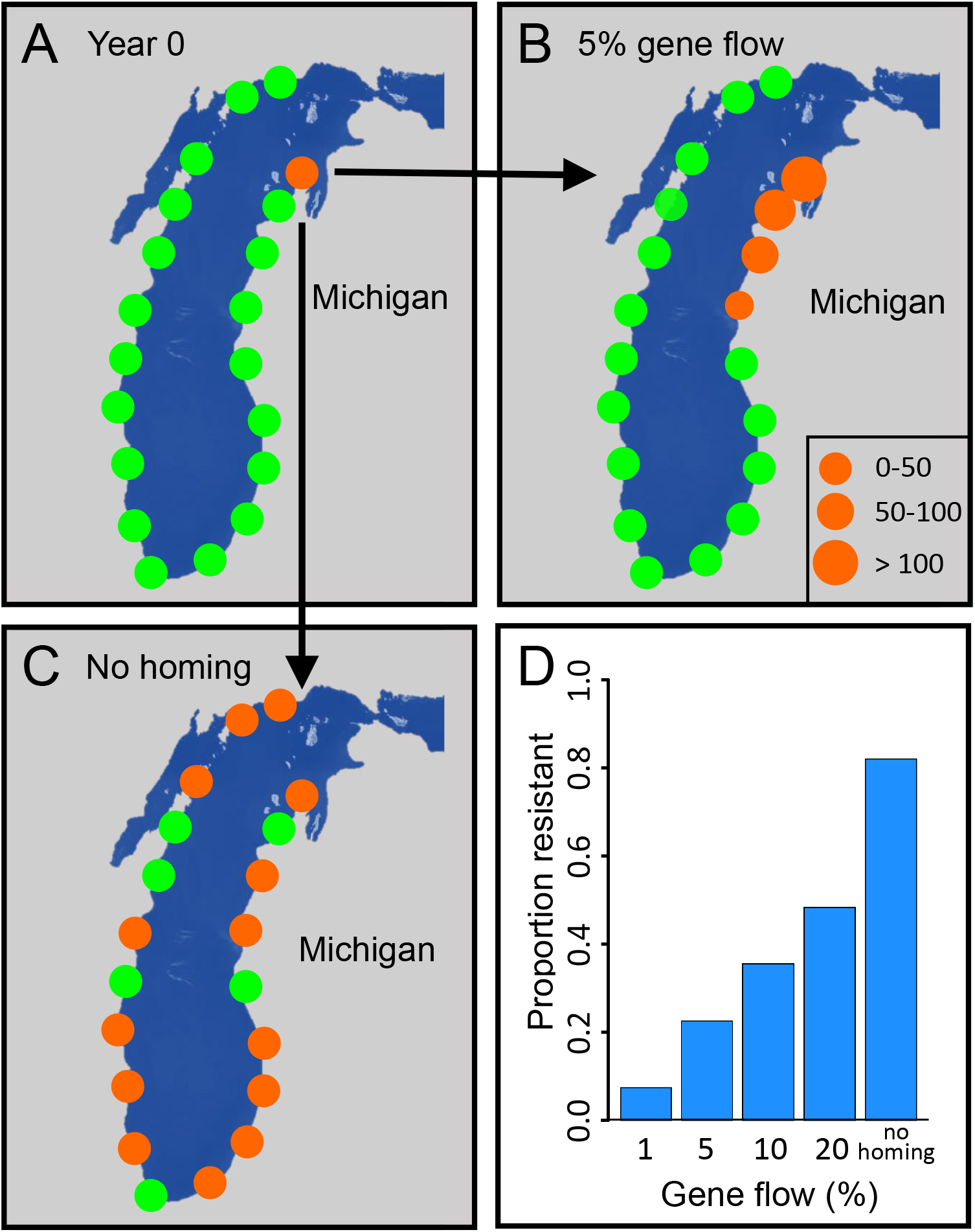
Spread of resistance for different dispersal and gene flow scenarios. A single resistant adult is introduced into one tributary of Lake Michigan (Panel A) where orange circles represent tributaries with at least one resistant larva and green circles represent tributaries with no resistant larvae. Assuming a stepping-stone pattern of isolation-by-distance and 5% gene flow of adults among populations, resistance spreads to three additional tributaries after 15 years (B). By contrast, when adults do not exhibit natal philopatry, as is commonly documented in sea lamprey populations, resistant lamprey spread to 70% of tributaries (C). Out of 100 replicates, resistance developed 82% of the time when there is no natal philopatry (here denoted as “no homing”), but develops substantially less often for lower rates of migration and gene flow (D). These results illustrate that once resistance develops, sea lamprey dispersal behaviors will result in the rapid spread of resistance throughout the system.

The lack of natal philopatry also makes the early detection of resistance challenging. Assuming that a resistant larva could be detected in a given sample, an immense sampling and screening effort is required to detect resistance in its early stages (e.g., 10 or 20 years after the introduction of a single resistant adult) (Figure 5). For example, if 0.5% of all larvae were sampled 20 years after the release of a single resistant adult, the probability of the sample including a single resistant larva would only equal 50%. Given that larval abundances are on the order of tens millions of individuals, a 0.5% sample size would involve screening a minimum of fifty thousand individuals. Furthermore, the probability of detection decreases as the costs of resistance increase (Figure S3).

**Figure 5.**
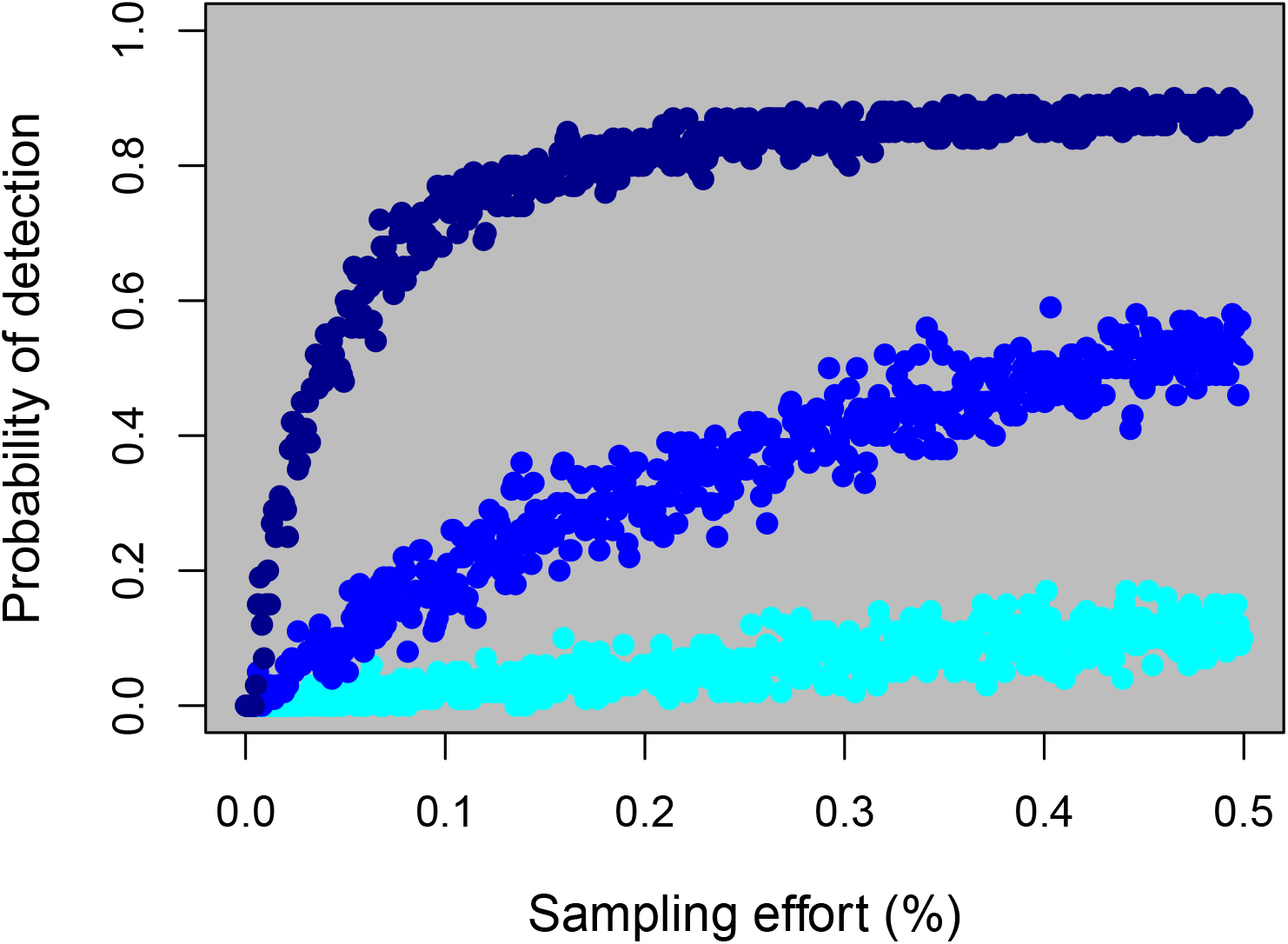
Relationship between the probability of detecting (i.e., sampling) a resistant individual for various sampling efforts (% of all larvae sampled), assuming no costs of resistance and perfect identification of resistant individuals. Sampling effort was conducted 10 years (cyan points), 20 years (blue points), and 30 years (dark blue points) after the introduction of a resistant adult. Larvae were sampled randomly across all tributaries and a total of 500 independent samples per time point were collected. Because the larval populations number in the tens of millions, these results illustrate that it can be challenging to detect resistance in its early stages without screening vast numbers of larvae.

We next examined the costs of resistance in this system by penalizing the fecundity and thus the reproductive success of resistant adult lamprey. To examine the consequences on larval abundance, we stopped TFM treatment at year 100. We found that when there was no cost of resistance, the total number of resistant larvae remained relatively constant for the next 100 years (Figure 6*a*). From a pragmatic standpoint, this result means that if there is no cost of resistance, then resistant genotypes cannot be easily purged from the system even in the absence of treatment. If there is a moderate cost to resistance (i.e., 10%), then the number of resistant larvae will decline through time in the absence of treatment, but not in a way that is meaningful from a management perspective (i.e., it can take 100 years or more for all resistant larvae to be eliminated from the system) (Figure 6*b*). By contrast, if there is a high cost to resistance (40%), then stopping treatment could be an effective management strategy and resistant larvae can be eliminated from the system in as little as 10 years (Figure S4).

**Figure 6.**
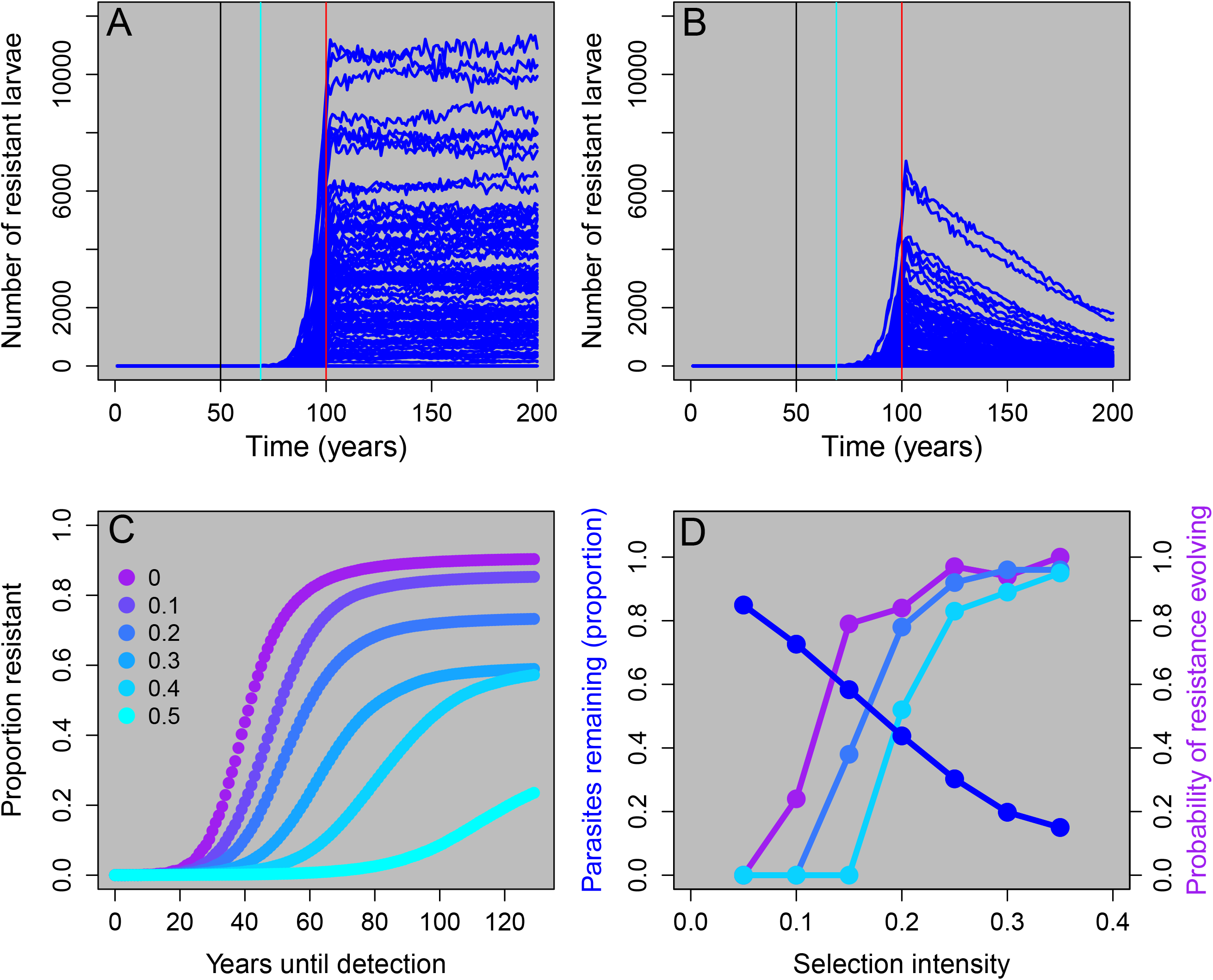
Effects of costs of resistance on the evolution of resistance. Panels A and B illustrate the relationship between the cost of resistance and the number of resistant larvae through time. TFM treatment was started in year 50 (black vertical line), a single resistant adult was added in year 70 (blue vertical line) and TFM treatment was stopped in year 100 (red vertical line). When there is no cost of resistance (panel A), the number of resistant larvae does not decline through time even after TFM treatment is stopped. When cost of resistance is moderate (10% reduction in fitness, panel B), the number of resistant larvae gradually declines through time. Panel C illustrates the proportion of resistant larvae as a function of years until detection (e.g., 0 equals model year 70). Cost of resistance was varied from 0 to 0.5 in increments of 0.1 (see legend). Notice that as the cost of resistance increases the total number of resistant larvae is smaller and the time for resistance to develop is longer. Panel D illustrates the trade-off between the proportion of parasites killed by TFM (measured as proportion of parasites remaining in the system immediately before the introduction of a resistant individual, blue line) and the probability of resistance evolving (defined as the proportion of 100 replicates with > 90% of larvae being resistant at year 200). To vary the selection intensity, we varied the number of streams treated each year and examined three costs of resistance where colors match the legend in panel C (no costs of resistance (purple line), a 20% cost of resistance, and a 40% cost of resistance. A higher cost of resistance shifts the probability of resistance evolving curve to the right, but notice that as selection intensity increases the probability of resistance evolving also increases.

We also examined how costs of resistance influence both the probability and the time it takes for resistance to develop in the system. We found that as the cost of resistance increases, the proportion of resistant larvae decreases (Figure 6*c*). We next illustrate a strong tradeoff between the proportion of parasites killed by TFM and the probability of resistance evolving (Figure 6*d*). As selection intensity increases, the proportion of the parasite population that is initially reduced increases, but so does the probability of resistance evolving (Figure 6*d*). As the cost of resistance increases, the probability of resistance evolving decreases, but only up to a point (see also Figure S5). Thus, costs of resistance can reduce, but not eliminate, both the likelihood and speed at which resistance develops in this system.

## Discussion

Our results indicate that sea lamprey, an invasive fish species successfully treated with a pesticide for over 60 years in the Great Lakes, could evolve resistance between 40 and 80 years after the introduction of a resistant individual. The single largest determinant of whether resistance takes a comparatively short (e.g., 30 years) or longer time (e.g., 80 years) to develop is the annual number of tributaries treated. Across all Great Lakes, sea lamprey have been treated with TFM for an average of 50 years^34^, which is towards the beginning of when our model predicts resistance will evolve. As such, sea lamprey in the Great Lakes could be in the incipient stages for the evolution of resistance. If alternative control measures can’t be found, the spread of resistance in Great Lakes sea lamprey could threaten the viability of one of the world’s most successful invasive vertebrate control programs and cause substantial declines in economically valuable commercial and recreational fish populations. Compared to most other systems studied to date, resistance in the lamprey-TFM system can be viewed as taking a relatively long time to develop (as measured in years, not generations), but rapidly accelerates after a certain threshold is reached (Figure 2). The relatively long time it takes for resistance to develop is because: *i*. sea lamprey can spend up to seven years in the larval stage before transforming into a parasitic adult and *ii*. the lack of natal philopatry results in resistant lamprey remaining at low abundance while spreading throughout the entire system. These life history characteristics mean that resistant larvae and adults initially increase in abundance very slowly in the system.

This slow initial accumulation of resistant individuals also means that it can be very challenging to detect resistance in its incipient stages. In fact, it can take 40 years of more before the proportion of resistant larvae increases above 50% (Figure 6*c*). Ten years after the introduction of a resistant adult, our model predicts that, for a moderate selection intensity, screening 0.5% of the larval population for resistance would only result in a 10% probability of detection, meaning that there would only be a 10% probability of including a single resistant individual in the sample. Because sea lamprey are highly fecund, there can be tens of millions of larvae in the population at any given point in time (GLFC control board, personal communication). Thus, a screening procedure that sampled a minimum of 50,000 larvae would only have around a 10% probability of having at least one resistant larvae 10 years after the introduction of a resistant adult (Figure 5). Our results also assume perfect detection; we assume that if a resistant individual was included in the sample then it would be correctly identified as such. In reality, type I errors (classifying a non-resistant individual as resistant) and type II errors (failing to detect a resistant individual when it is present in the sample) would both affect the probability of successful detection. Technical issues aside, the fact remains that a very large sample of larvae must be screened to detect resistance during its early stages. This detection challenge presents a quandary; how much effort and resources should be devoted to screening for resistance versus developing new control measures?

Including a fitness cost for resistance decreased the time for resistance to be eradicated from the system via natural selection once control with TFM ceased. However, resistant individuals were eliminated from the system within a short enough time frame acceptable for management (e.g., 20 years or less) only if the cost of resistance was very high (0.4). We further found that the onset of resistance is delayed and the total number of simulations that developed resistance decreased with increasing costs of resistance. However, regardless of the costs of resistance, there was a strong trade-off between the proportion of parasites initially controlled by TFM and the probability of resistance evolving (Figure 6*d*). The relative roles of density dependence and density independence in evolution has often been ignored (but see ^48,49^). Here, we show that as the relative strength of density dependent reproduction decreases, the proportion of replicate populations that develop resistance decreases and, for those that do develop resistance, the number of years until resistance develops increases (Figure S6). Although, it is believed that sea lamprey in the Great Lakes are highly regulated by compensatory dynamics^44^, robust empirical data regarding density-dependent dynamics during reproduction and shortly thereafter are largely lacking. Thus, both high costs of resistance and reduced density dependent reproduction can delay, but not entirely prevent, the evolution of resistance.

### Caveats, assumptions, and management recommendations

Our model results suggest that the onset of resistance is either ongoing or imminent for invasive Great Lakes sea lamprey that have been treated with TFM for over 60 years. There are, however, a few caveats worth considering. First, it is possible that the total percentage of larval habitat treated with TFM each year is overestimated. Adults may spawn in undocumented areas and there could be large areas of larval habitat that go unsurveyed (e.g., small reaches of tributaries, lentic bays). If the percentage of treated larval habitat is smaller than expected, the overall mortality rate imposed on the population by TFM could be lower than expected which would reduce the selection intensity (Figures 2,6). However, the ability of the control program to keep adult sea lamprey numbers at relatively low levels implies a mortality rate high enough to impose some selective pressure. Second, we have assumed that resistance will evolve via genes of large effect, the most common form of resistance documented in the literature^50,51^. However, it is possible that resistance could evolve more along the lines of a quantitative trait with hundreds of loci. If this were to happen, then the response to selection is expected to be much slower^52^, and the onset of resistance could be delayed by tens of generations. Of course, resistance can evolve in multiple ways and at any point in time, especially given the lamprey’s high fecundity, and so the evolution of a different class of resistant individuals with a gene of large effect may be more likely to occur before the quantitative trait resistance alleles increase in frequency. Third, we don’t consider the effects of prey abundance (or other factors like climate) on the sea lamprey population size. Given that sea lamprey population sizes do not affect the timing or magnitude of resistance (Figures S1, S2), we do not expect to see much of an effect for prey availability on resistance evolution. Nonetheless, small effective populations may limit the response to selection (but see Wood, et al. ^53^ or there could be unaccounted eco-evolutionary feedbacks such as how energy reserves affect fecundity or timing of reproduction . Lastly, it is possible that a severe founder effect or a unique mode of action for TFM may limit the likelihood of an evolutionary response. We think these possibilities are unlikely. Microsatellite data suggest that genetic diversity in Great Lakes sea lamprey is not substantially lower than lamprey in their native range^38^. Furthermore, because the mode of action for TFM largely targets ATP synthesis^54^ there are many potential modes of resistance evolution^34^.

Considering these results, we have several management recommendations. First, we suggest that alternative control methods with different modes of action and high target specificity be developed as quickly as possible. In the meantime, one option is to reduce the strength of selection by lowering the number of treated streams. This decision would have to be carefully considered against the shorter-term economic and ecological costs of potentially higher numbers of parasites. Through a risk assessment and cost-benefits analysis, it should be possible to identify a target level of streams to treat that maximizes the tradeoff between parasite abundance and resistance evolution (e.g., Figure 3). Second, it would be prudent to develop a high throughput screening program aimed at detecting resistant larvae. Finally, there are several important research gaps that, if addressed, would enable a more accurate assessment of the timelines of resistance evolution in this system and enable more effective management^55^; these include quantifying the degree of density-dependent reproduction, characterizing fine-scale genetic spatial structure, measuring the true strength of selection, and studying potential costs of resistance. Appropriate action to impede the onset of this incipient crisis should remain a high priority for the management and conservation of Great Lakes fishes. Such action could not only prevent the loss of billions of dollars from the economy but could also prevent large-scale and undesirable ecological shifts in abundance and community composition.

## Data availability

Model code and scripts available at https://github.com/ChristieLab

## Acknowledgements

We thank A Martinez, A Muir, C Searle, and M Siefkes for feedback and assistance with this project. This research was facilitated by the Rosen Center for Advanced Computing at Purdue, West Lafayette, IN. This project was funded by support to MRC and MSS from the Great Lakes Fishery Commission (Project ID: 2016_CHR_54053).

## Author contributions

M.R.C, M.S.S., and E.S.D. developed the conceptual framework, guided model development, and wrote the manuscript. M.R.C. wrote the model and analyzed the data.

