## Supplemental Information for "Rapid resistance to pesticide control is predicted to evolve in an invasive fish"

**Supporting Information:** Invasive sea lamprey may soon become resistant to their primary control agent, 3-trifluoromethyl-4-nitrophenol

**Supporting Methods:**

*Sea lamprey life history:*

Throughout its native range, which includes much of the Northern Atlantic, the anadromous sea lamprey has a bipartite life cycle consisting of a filter-feeding larval stage and a parasitic adult stage. The larval sea lamprey, known as ammocoetes, remain buried in detritus-rich substrate where they filter feed for an average of 5 to 8 years (documented range 2 to 19 years) (Renaud 2011). The metamorphosis of larval sea lamprey into adults involves substantial behavioral and physiological modifications required for their hematophagous, parasitic lifestyle (Youson 2003). After metamorphosis, juvenile sea lamprey enter the ocean and begin searching for hosts. Sea lamprey have been observed feeding on a diverse array of species such as herring, mackerel, and salmon (Kottelat & Freyhof 2007). Sea lamprey attach to their hosts with rows of sharp teeth and they secrete anticoagulants that allow them to feed continuously (Gage & Gage-Day 1927). Throughout their native range, sea lamprey typically do not kill their hosts and an individual lamprey will often switch between hosts (Kottelat & Freyhof 2007). After feeding for 20 to 36 months, the lamprey mature and begin to search for rivers and streams suitable for spawning, migrating anywhere between 20 to 850 km inland. Unlike other anadromous fishes, such as salmon, sea lamprey do not exhibit natal philopatry. Instead, sea lamprey cue in on chemicals produced by larvae living in freshwater streams and rivers (Bjerselius et al. 2000; Sorensen & Hoye 2007). This reproductive strategy means that sea lamprey populations are largely panmictic, though there is evidence for restricted gene flow between populations on opposite sides of the Atlantic Ocean (Bryan et al. 2005; Waldman et al. 2008). Sea lamprey are strictly semelparous; after spawning all adults die.

In the Great Lakes, sea lamprey retain most of these life history characteristics, however, there are some notable exceptions. First, metamorphosed sea lamprey treat the Great Lakes as a surrogate ocean, thus they never physiologically acclimate to a salt-water environment. Second, growth rates are faster, the larval duration is shorter, adults are smaller, and fecundity is slightly lower (Heinrich et al. 1980; Young et al. 1990). Lastly, sea lamprey in the Great Lakes spend a longer period attached to a single host and an individual host can have many more sea lamprey attached to them than host species found in the Atlantic Ocean. This last difference, possibly due to higher adult lamprey abundance and/or a greater lamprey to host ratio, means that sea lamprey are directly responsible for mortally wounding large numbers of host fishes in the Great Lakes (Farmer & Beamish 1973). Even if sea lamprey do not directly kill their hosts, many host fish later die from subsequent fungal infections associated with lamprey parasitism and fish that continue to survive have been shown to have decreased reproductive success (Swink & Hanson 1989; Swink 1990).

*Density-dependent reproduction:*

We next varied the strength of density-dependent reproduction by varying the parameter $n_{m}$ in equation 3. We varied this parameter to have values ranging from 50 to 2 resulting in a relative strength of density-dependent reproduction ranging from 1 to 0.04 while using the default values for all other parameters (Table 1). We measured both the proportion of resistant populations (defined as the proportion of 100 replicates with > 90% of larvae being resistant at year 200) and the number of years until resistance evolved (defined as the average number of years each of the replicates (excluding replicates with no resistant larvae) took until > 90% of larvae were resistant) as a function of the relative strength of density dependent reproduction. We also varied the magnitude of density independent mortality (as opposed to reproduction) for both larvae and juveniles, though these parameters had little effect on the evolution of resistance.

S**upporting Results:**

*Density dependent reproduction:*

Lastly, we examined the role of population regulation on the evolution of resistance by varying the strength of density-dependent reproduction. As we increased the relative strength of density-independent reproduction (i.e., number of offspring surviving their first few months), we found that the proportion of resistant populations found at year 200 decreased substantially (figure S6). In fact, when there is only density-independent reproduction (i.e., no density dependence) the eventual evolution of resistance becomes very rare. We also found that as density independence increased the average number of years until resistance developed initially decreased, but then increased substantially. Interestingly, the magnitude of density independent survival for both larvae and juveniles had almost no effect on these parameters such that the stage at which density dependence or independence predominates may be more important than their relative magnitudes (Baskett & Waples 2013).

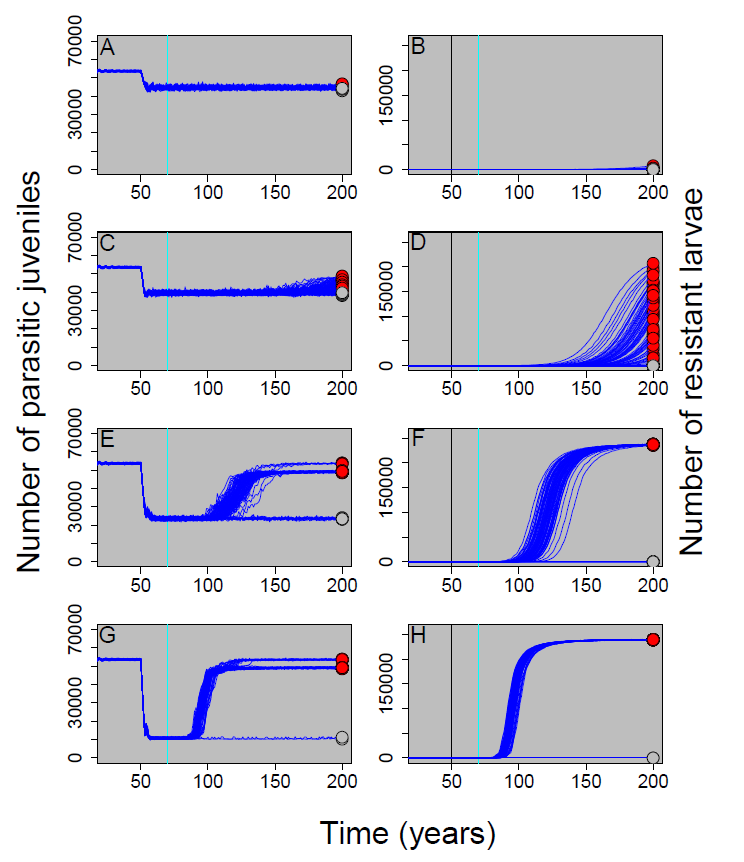

**Figure S1**. Parasitic juvenile and larval abundances through time where the proportion of tributaries treated each year equaled 0.05 (panels A, B), 0.1 (panels C, D), 0.2 (panels E, F), and 0.3 (panels G, H), respectively. TFM treatment was started at year 50 and a single resistant adult was introduced in year 70 (vertical blue line). One hundred replicates (dark blue lines) were run for each scenario and at year 200, red circles represent populations with resistant individuals and grey circles represent populations with no resistant individuals. As the number of tributaries treated each year increases there is a decrease in the number of parasitic individuals, but also a rapid increase in the number of resistant larvae. In comparison with the main text, these simulations were run with an order of magnitude higher larval (12000 vs 1200 per tributary) and juvenile carrying capacities (80000 vs 8000) and the qualitative patterns remain unchanged (*cf.* figure 2)

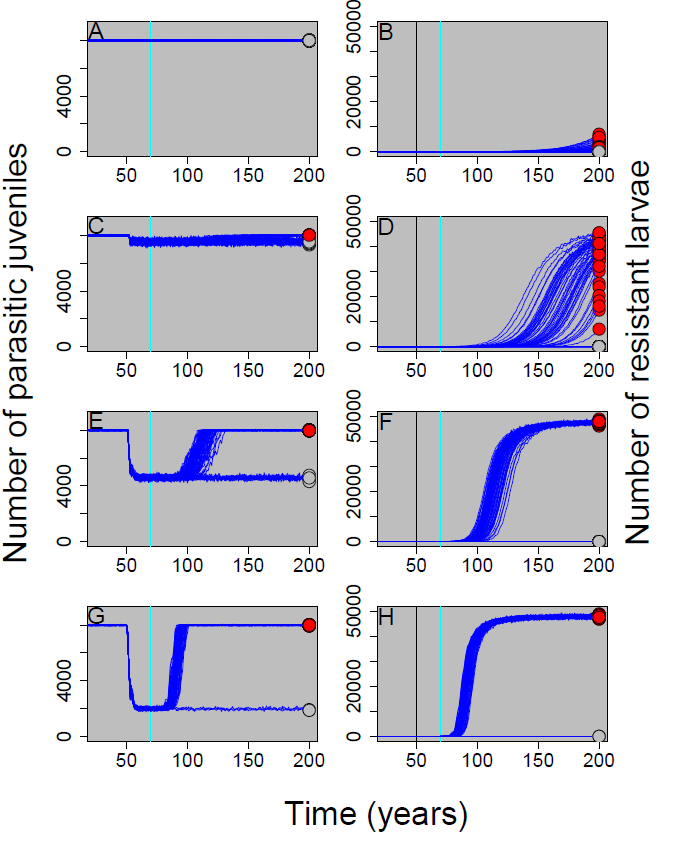

**Figure S2**: Parasitic juvenile and larval abundances through time where the proportion of tributaries treated each year equaled 0.05 (panels A, B), 0.1 (panels C, D), 0.2 (panels E, F), and 0.3 (panels G, H), respectively. TFM treatment was started at year 50 and a single resistant adult was introduced in year 70 (vertical blue line). One hundred replicates (dark blue lines) were run for each scenario and at year 200, red circles represent populations with resistant individuals and grey circles represent populations with no resistant individuals. As the number of tributaries treated each year increases there is a decrease in the number of parasitic individuals, but also a rapid increase in the number of resistant larvae. In comparison with the main text, these simulations were run with twice the number of tributaries as the default model (40 vs. 20) yet the qualitative patterns remain unchanged (*cf.* figure 2). Results for 60 tributaries were nearly identical to this figure and figure 2 (data not shown).

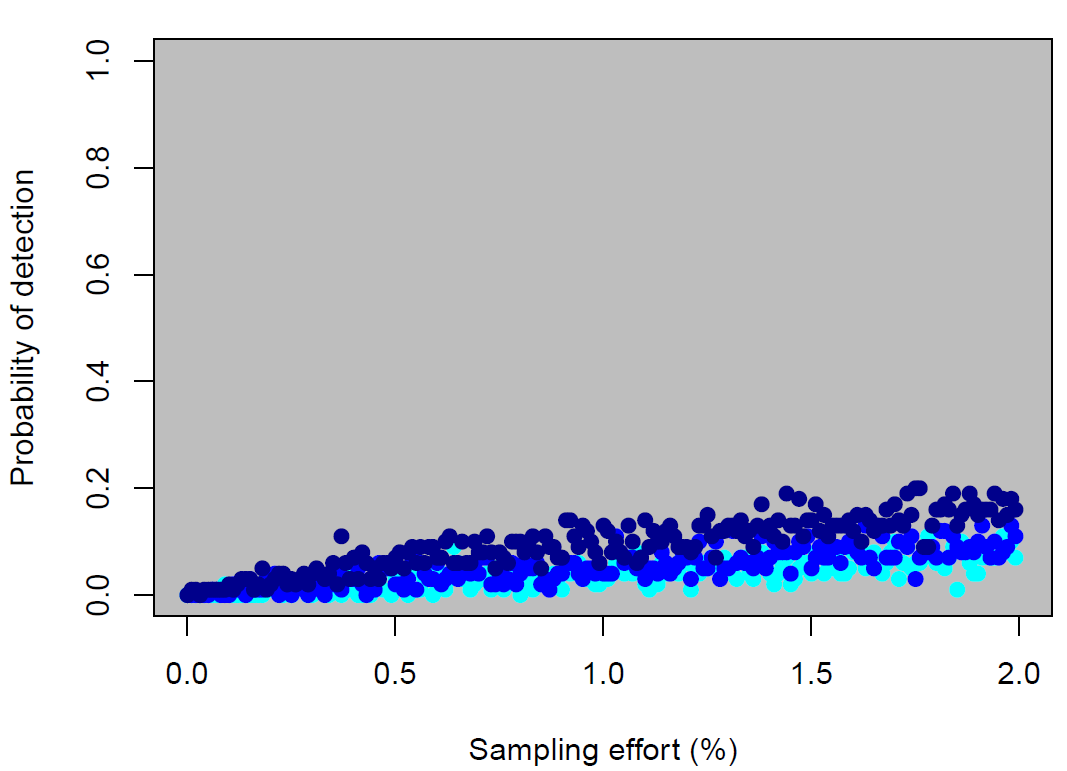

**Figure S3**: Assuming a 50% fitness cost of resistance and perfect identification of a resistant individual, this figure illustrates the probability of detecting (i.e., sampling) a resistant individual for various sampling efforts (% of all larvae sampled). Sampling effort was conducted 10 years (cyan points), 20 years (blue points), and 30 years (dark blue points) after the introduction of a resistant adult. These results illustrate that resistance can take a relatively long time to develop in this system and that it can be challenging to detect resistance in its early stages. Costs of resistance make detection even more challenging (compare with figure 5 in the main text where there are no costs of resistance).

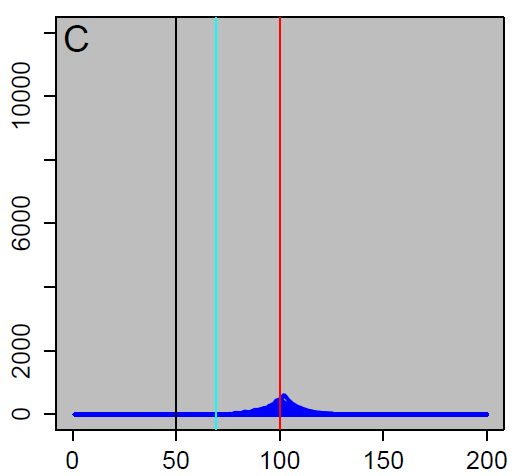

Number of resistant larvae

Time (years)

**Figure S4.** Relationship between the cost of resistance and the number of resistant larvae through time. TFM treatment was started in year 50 (black vertical line), a single resistant adult was added in year 70 (blue vertical line) and TFM treatment was stopped in year 100 (red vertical line). Individual simulation results for 100 replicates are shown. When the cost of resistance is high (40% reduction in fitness depicted here), there are smaller numbers of resistant larvae at year 100 and the resistant larvae are rapidly eliminated from the population. No costs or more moderate costs of resistance (figure 6 *A, B*), result in resistant larvae remaining in the system for much longer periods of time.

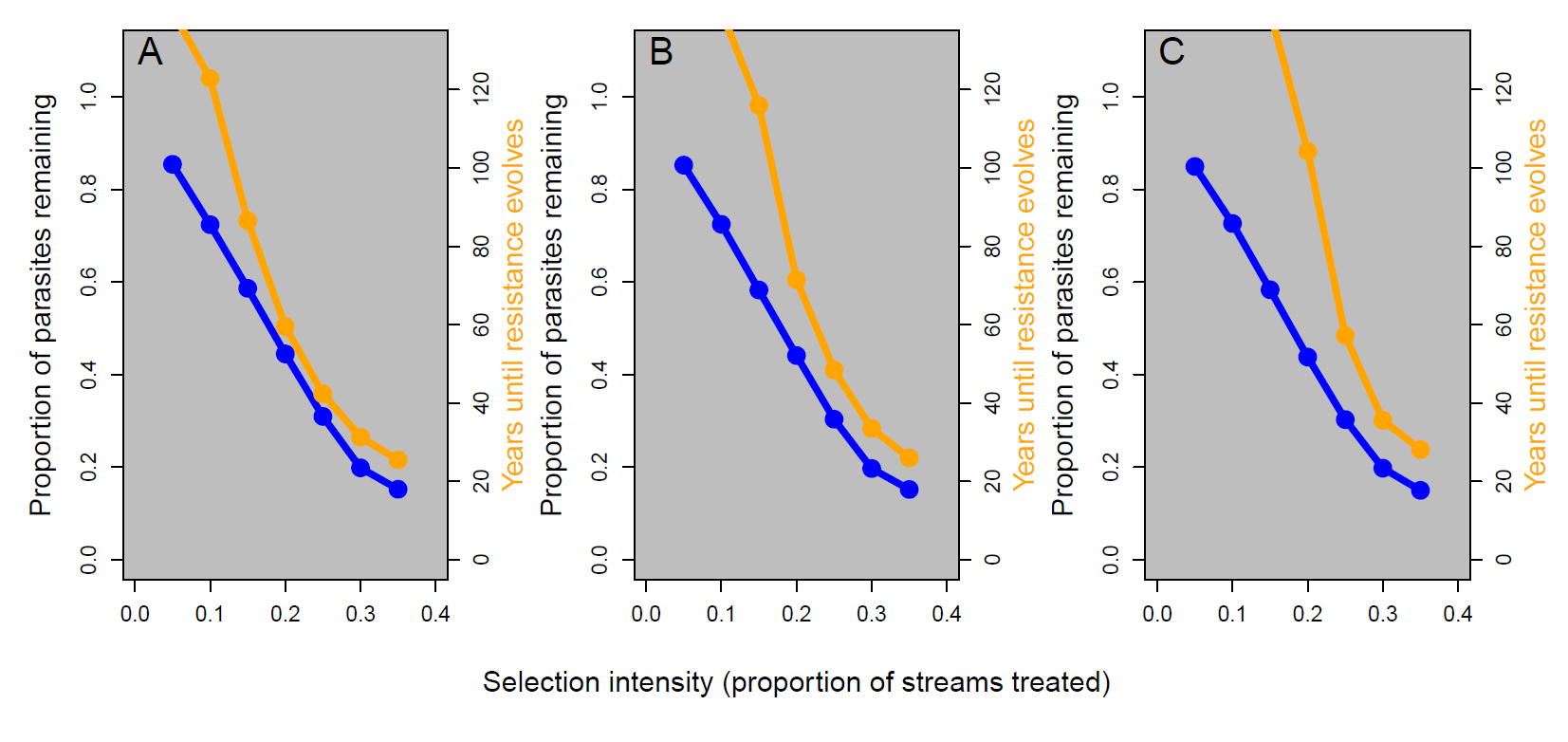

**Figure S5**: Relationship between the proportion of parasites killed by TFM (here measured as proportion of parasites remaining in the system before the spread of resistant individuals, blue lines) and the number of years until of resistance evolves (defined as the proportion of 100 replicates with > 90% of larvae being resistant at year 200, orange lines). To vary the strength of selection, we varied the number of streams treated each year and examined three costs of resistance where panel A equals no cost of resistance, panel B equals a 20% cost of resistance, and panel C a 40% cost of resistance. A higher cost of resistance shifts the time-until-resistance curve to the right but notice that in all cases that as the strength of selection increases (i.e., the more larvae killed by TFM each year) the time until resistance evolves decreases.

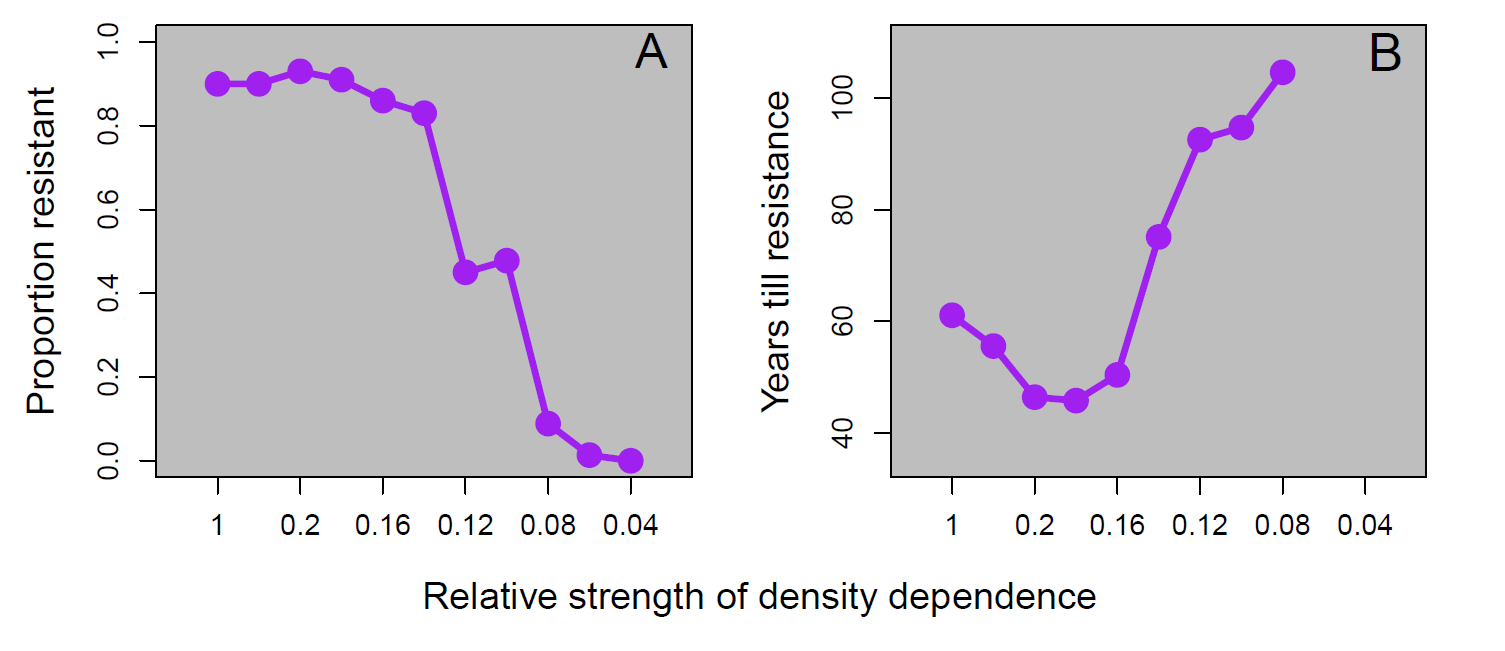

**Figure S6**: The proportion of simulations (out of 100) where resistance evolved and the number of years until resistance evolves (defined as the average number of years each of the replicates took until > 90% of larvae were resistant excluding replicates with no resistant larvae at year 200) as a function of the relative strength of density dependent reproduction. As the magnitude of density dependent reproduction decreases the proportion of resistant populations also decreases (A). By contrast, as the relative strength of density independence increases, the number of year until resistance develops initially decreases, but then increase substantially (B).
